# Microbial succession at weaning is guided by microbial metabolism of host glycans

**DOI:** 10.1101/2025.02.20.639370

**Authors:** Jean-Bernard Lubin, Paul J. Planet, Michael A. Silverman

**Author notes:** Michael A. Silverman, Paul J. Planet (. Co-senior authors (Paul J. Planet, Michael A. Silverman).

## Abstract

The weaning transition from a milk-based to a solid-food diet supports critical developmental changes to the intestinal microbiome and immune system. However, the specific microbial and host features that influence microbial succession at weaning are not well understood. Here, we developed a simple approach to investigate the complex dynamics of microbial succession during weaning by co-housing gnotobiotic mice colonized with the defined pre-weaning community PedsCom and the adult-derived consortium OMM12. As expected, co-housing PedsCom mice with OMM12 recapitulated microbial succession at weaning and induced immune system maturation in PedsCom mice. Unexpectedly, we found that the OMM12 microbes with the highest host glycan utilization profiles were the most adept colonizers of PedsCom mice. Genomic analysis confirmed that PedsCom is deficient in the carbohydrate-active enzymes responsible for degrading host-derived glycans, including mucins, compared to adult-derived consortia. We validated a role for glycan utilization *in vivo* by demonstrating that the mucus-degrading commensal microbe *Akkermansia muciniphila* critically depends on the metabolism of mucin glycans for colonization of PedsCom mice. These findings highlight the importance of host-derived glycans in shaping microbial communities during the weaning transition and suggest host glycans as novel targets to modulate intestinal microbial populations, introduce beneficial probiotics, and enhance immune system development during weaning.

## Introduction

The intestinal microbiomes of mammals are complex and dynamic communities that are profoundly influenced by the nutrients available in the gut^1,2^. Diet has a considerable impact on early-life microbiome composition, as demonstrated by the well-established differences in the microbiomes of infants fed breast milk versus formula, driven primarily by host-indigestible milk oligosaccharides (MOs)^3,4^. All mammals undergo weaning, the important transition from a solely milk-based diet to solid food. The weaning transition is associated with a rapid and dramatic increase in microbial diversity, changes in microbial composition, and corresponding immune system maturation^5^. During weaning, dietary carbohydrates shift from lactose and MOs found in milk to starches and host-indigestible fibers found in solid foods. The shift from MOs to fiber is thought to be the principal driver of microbial succession and community composition during the weaning transition^6,7^, yet, due to the complexity of gut microbiomes, there are limited experimental systems to identify the specific microbes and metabolic pathways that drive microbiome and immune system maturation at weaning.

We recently developed a nine-bacteria consortium called PedsCom to model the unique composition and function of the mammalian pre-weaning intestinal microbiome^8^. PedsCom mice microbiomes are locked into a pre-weaning configuration, which leads to stunted immune system maturation and high susceptibility to enteric infection^8^. The PedsCom consortium has a metabolic profile distinct from gnotobiotic communities derived from adult mice^9^. In particular, the PedsCom consortium is enriched in genes that encode enzymes that degrade lipids and amino acids, nutrients that are abundant in milk. Here, we developed a unique, tractable *in vivo* system to experimentally model the complex dynamics of microbial succession from a pre-weaning to an adult microbial community by co-housing PedsCom mice with an adult-derived gnotobiotic community, Oligo-Mouse-Microbiota 12 (OMM12)^10^. Unexpectedly, we found that the microbial capability to utilize host glycans is the strongest predictor of which adult-associated microbes will colonize PedsCom mice. We validated the *in vivo* role of microbial utilization of host glycans by demonstrating that metabolism of mucus-derived glycans allows *Akkermansia muciniphila* to colonize ~100-fold higher abundance in PedsCom mice.

## Results

### Oligo-MM12 strains ‘mature’ PedsCom mice

The OMM12 and PedsCom consortia model the metabolic functions of complete adult and pre-weaning intestinal microbiomes, respectively^10^. To investigate microbial succession of the weaning transition using defined consortia of microbes, we co-housed adult OMM12 mice with adult PedsCom mice for two days to allow horizontal transfer of microbes, removed and sacrificed the OMM12 mice, and then allowed the intestinal communities of co-housed PedsCom recipient mice to stabilize for five weeks (**Fig. 1A**). We predicted that bacteria from OMM12 that are better suited to the adult diet would preferentially colonize PedsCom mice. Ribosomal RNA gene 16S-based sequencing of small intestinal, cecal and colonic samples in co-housed PedsCom mice revealed almost complete replacement by OMM12 (~90%) strains across the length of the gastrointestinal (G.I.) tract (**Fig. 1B**). This pattern of succession dominated by adult-associated microbes mirrors the changes in microbiome composition during the natural process of weaning in mice and humans^2,11^.

**Figure 1.**
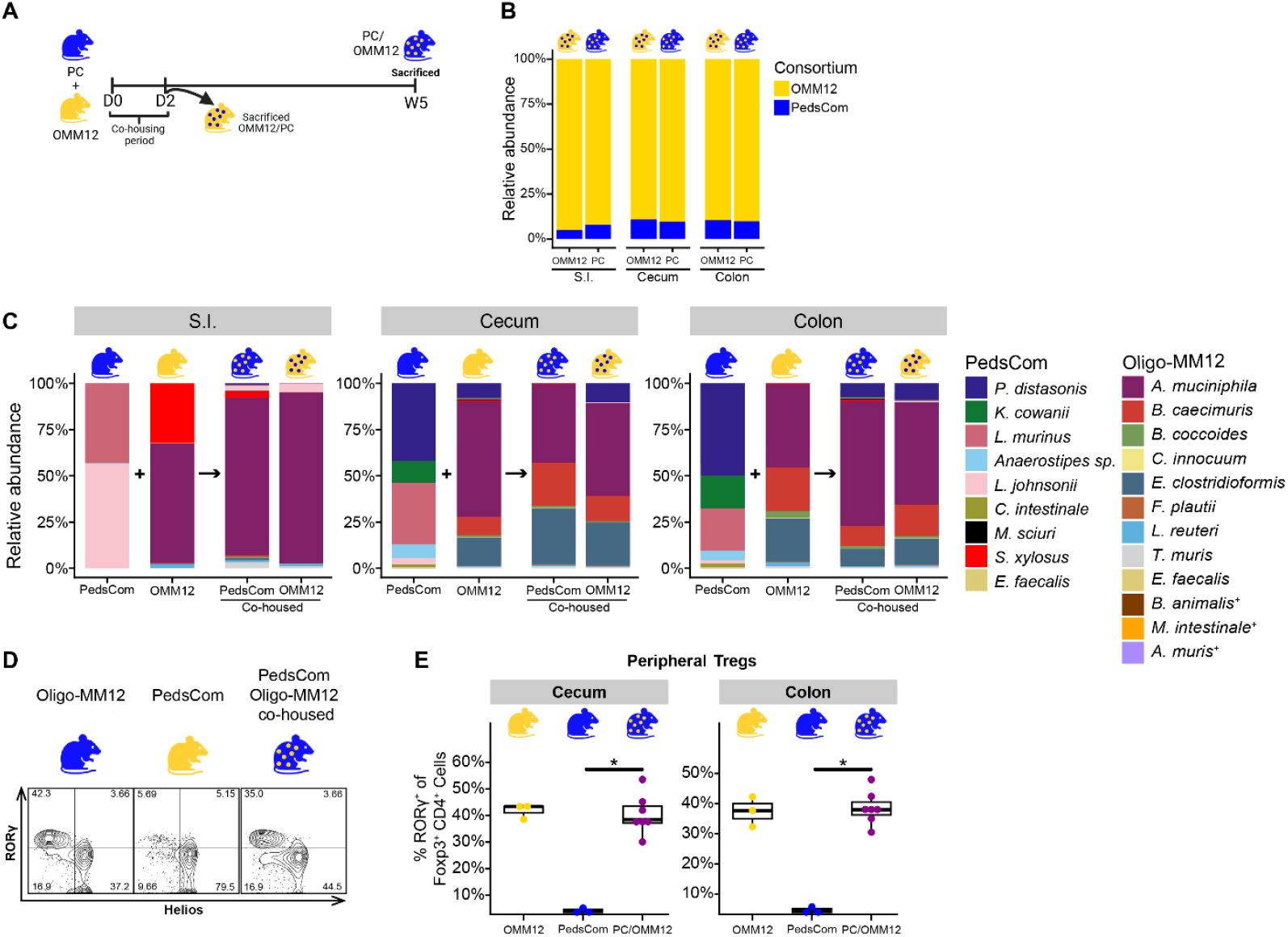
Oligo-MM12 strains ‘mature’ PedsCom mice. **A**. Schematic of PedsCom and Oligo-MM12 co-housing experiment. **B**. 16S rRNA gene sequencing relative abundances of consortia in Oligo-MM12 (N = 2) and PedsCom (PC) mice (N = 7) post co-housing. **C**. Taxa bar plots of the mean relative abundances of PedsCom and Oligo-MM12 strains in the small intestine, cecum and colon by 16S rRNA gene sequencing during the co-housing experiment. Representative data from control Oligo-MM12 and PedsCom mice is provided for comparison. **D**. Representative flow plots of the gating scheme for immune cells population analysis of large intestine and cecum. **E**. Peripheral regulatory T cells populations of Oligo-MM12 [(OMM12) N = 3] and PedsCom [(PC) N = 3] control mice, and PedsCom co-housed mice [(PC/OMM12) N = 7] in the cecum and colon. Dot plots are shown with median value. Mann-Whitney-Wilcoxon tests performed, *p<0.05. + = Strains that were undetectable in Oligo-MM12 colonized mice.

*Akkermansia muciniphila* was the most abundant colonizer of PedsCom co-housed mice, encompassing 85% of the small intestine and >60% of the cecum and colon communities (**Fig. 1C and S1A**). *Bacteroides caecimuris* and *Enterocloster clostridioformis* also readily colonized PedsCom mice. As expected, PedsCom strains competed poorly against the OMM12 microbes, with 5 out of 9 PedsCom members becoming undetectable following co-housing (**Fig. 1C**). The most persistent PedsCom members were *Parabacteroides distasonis*, primarily in the large intestine (7.6%), and *Lactobacillus johnsonii* in the small intestine (2.6%) (**Fig. S1B**). The drastic reduction in PedsCom species following OMM12 exposure supports the established paradigm that solid food at weaning provides nutrient sources and microbial niches for adult-associated microbes that are less accessible to early-life intestinal microbiota, but the specific nutrients and metabolic pathways that drive weaning-associated succession remain unclear.

Since critical immune cell populations are induced around weaning^12^, and adult PedsCom mice exhibit stunted immune development relative to mice with mature microbiomes^8^, we investigated whether the succession of adult-associated microbes into adult PedsCom mice would restore key defects in immune system maturation. We found that colonizing PedsCom mice with OMM12 strains significantly increased the canonical weaning-associated peripheral regulatory T cell (pTregs) populations^13^ in the cecum and colon of PedsCom mice (**Fig. 1D-E**). Since weaning-associated bacteria induce intestinal pTregs, we conclude that succession with adult-derived OMM12 microbes supports maturation of the immune system of PedsCom mice.

In summary, introducing adult-associated microbes from OMM12-colonized mice to PedsCom-colonized mice leads to microbiome succession to an adult-like microbiome configuration and immune system maturation in the PedsCom mice. This novel approach provides an experimentally tractable *in vivo* system to investigate the microbial features that drive microbial succession at weaning.

### Microbial metabolism of host-derived glycan predicts successful colonization of PedsCom mice

Weaning introduces major changes to the microbially-available carbohydrates in the intestine^6^. Since carbohydrate utilization profiles of gut commensals provide insight into niche specialization and microbial succession^14^, we compared carbohydrate metabolism of PedsCom to OMM12 using the predicted proteomes from the genomes of each strain and the Carbohydrate-Active enZYmes Database (CAZy) database^15^, which describes the families of structurally-related catalytic and carbohydrate-binding modules (or functional domains) of enzymes that degrade, modify, or create glycosidic bonds. Briefly, predicted protein sequences of each strain were analyzed using the dbCAN4 annotation tool^16^, and for each consortium we developed a CAZyme profile composed of proteins predicted to cleave, generate, or modify glycoside bonds. CAZymes were assigned to CAZyme families based on structural features and functional similarities. To assess the generalizability of potential differences between PedsCom and OMM12 strains, we included three additional adult-derived consortia: Altered-Schaedler Flora (ASF), GM15, and Oligo-Mouse-Microbiota 19 (OMM19)^17–19^. A CAZyme family was considered present in a consortium if it was identified in at least one strain. We manually annotated CAZyme families by target substrate (fiber, starch and oligosaccharides, host glycans, bacterial glycans, fungal glycans, or mixed function) to generate non-redundant CAZyme profiles of each community. A total of 358 CAZyme families of known function were annotated across the 47 unique strains from these four consortia. PedsCom contained fewer CAZymes per strain (92) than adult-derived consortia (OMM12 = 105, ASF = 117, GM15 = 133, OMM19 = 111) (**Fig. 2A**). This lower predicted CAZyme functionality in PedsCom is particularly illustrated by comparing to ASF, which encodes 100 more CAZymes than PedsCom despite being a smaller community (**Fig. S2A**). A comparison of the CAZyme family profile of each consortium offered insight into the diversity of putative carbohydrate usage of each community. PedsCom contains far fewer CAZyme families (156) than GM15 (265), OMM12 (225) and OMM19 (287), and there are significant differences in the substrate utilization profiles (**Fig. 2B**). PedsCom contains a comparable number of CAZyme families to ASF (156 v. 161), despite a higher absolute count of individual CAZymes detected in ASF. Overall, this *in silico* analysis provides evidence that PedsCom has significantly fewer CAZymes and a different carbohydrate usage profile than three rationally-designed adult consortia. The lower number of CAZymes suggested that PedsCom has a distinct and more focused carbohydrate utilization profile than the adult-associated communities.

**Figure 2.**
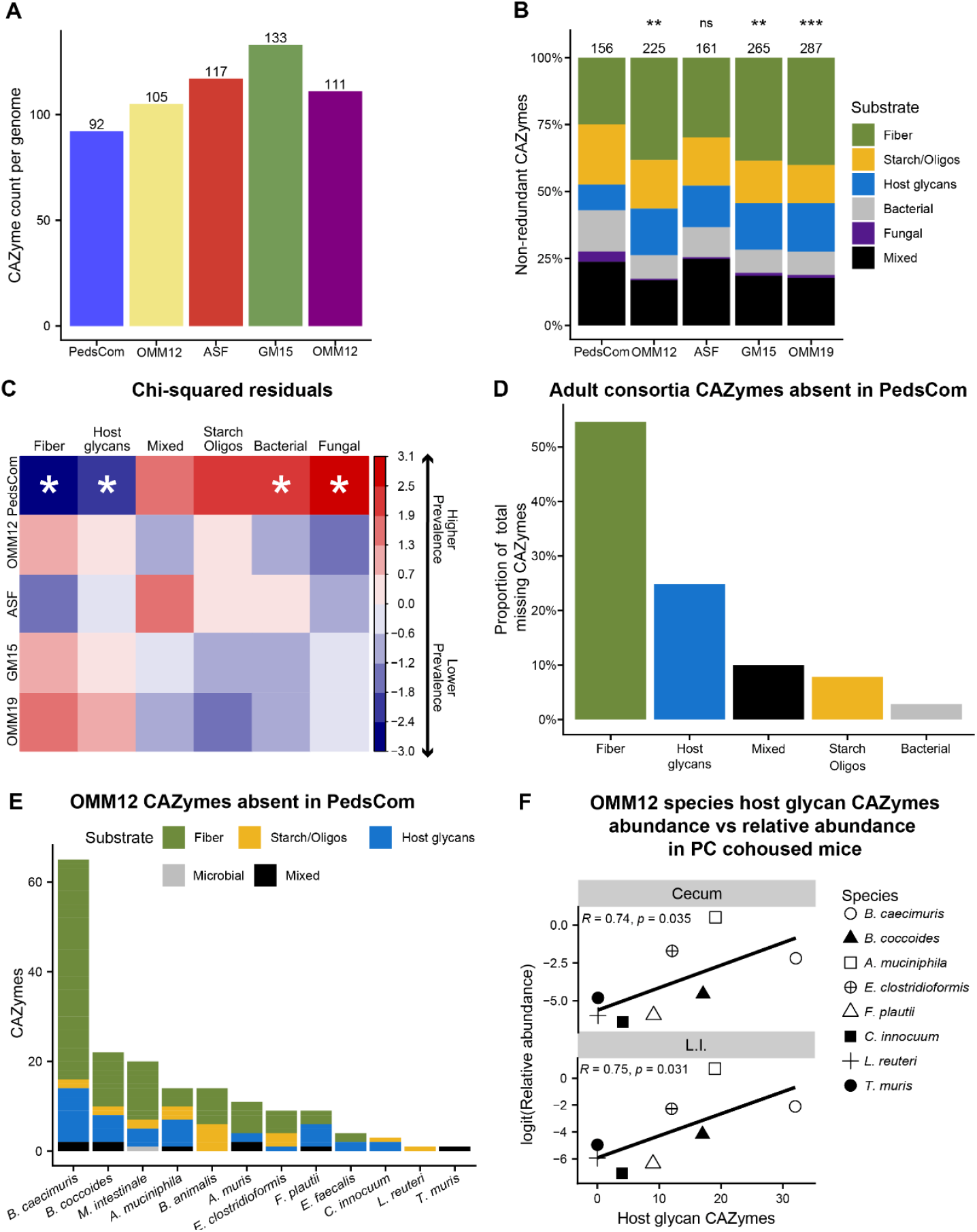
Microbial metabolism of host-derived glycan predicts successful colonization of PedsCom mice. **A**. Bar plot of annotated CAZymes in each consortium normalized by the number of member strains. Consortium: PedsCom (PC) = 9, Altered Schaedler Flora (ASF) = 8, GM15 = 15, Oligo-MM12 (OMM12) = 12, Oligo-MM19 (OMM19) = 19. **B**. Stacked bar plot of CAZyme families found in each consortium stratified by target substrate. Chi-squared test comparing the proportions of CAZyme families by target substrate found PedsCom to each adult-derived consortia, **p<0.01, ***p<0.001. **C**. Pearson residuals of Chi-square test between all consortia (*p = 0.01). Residual magnitude [increased (+) or decreased prevalence (-)] represents the effect size of the respective category in the significant difference between the consortia. **D**. Bar plot of the distribution of CAZymes not found in PedsCom but present in at least two adult-derived consortia by target substrate. **E**. Stacked bar plot of Oligo-MM12 strain CAZyme families missing in PedsCom. Strains are ordered by the number of CAZyme families and are colored by target substrate. **F**. Spearman correlations of host glycan CAZyme abundance in Oligo-MM12 strains and their relative abundances in PedsCom co-housed mice.

Due to a paucity of fiber in the pre-weaning milk-based diet, we hypothesized that PedsCom would encode fewer CAZymes associated with fiber metabolism than adult-derived consortia. Chi-squared analysis revealed a significant relationship between the PedsCom and the adult consortia and CAZyme family proportions (**Fig. 2B**). As expected, fiber-degrading CAZyme families provide the largest differences between the consortia. Unexpectedly, PedsCom also has a lower proportion of CAZyme families capable of degrading host-derived glycans (9.6% vs 14.6%, PC vs OMM12) (**Fig. 2B-C**). The missing host glycan families were primarily associated with degrading the carbohydrate components of mucins (sialidases, α-fucosidases, and α-*N*-acetylglucosaminidases) and glycosaminoglycans (chondroitin and hyaluronate lyases). Accordingly, most of the CAZyme families present in the adult-derived consortia, but absent in PedsCom, were associated with either fiber or host glycan metabolism (**Fig. 2D**). The lower number of CAZymes detected in PedsCom and the decreased prevalence of fiber-associated CAZyme families is consistent with the limited diversity of carbohydrate substrates in a milk-based diet^20–22^. Most intriguingly, the decreased prevalence of CAZyme families that degrade host-derived glycans and fiber in PedsCom microbes suggests that pre-weaning microbiota are adapted to thrive in an environment with fewer host glycan and fiber substrates.

Given the paucity of fiber and host-glycan degrading CAZymes in PedsCom, we considered the possibility that the adult-derived species that most successfully colonized PedsCom in the co-housing experiment (**Fig. 1A**) would simply be those that had the highest number of CAZymes missing in PedsCom. *Bacteroides caecimuris* contains the highest number of CAZymes missing in PedsCom (65), followed by *Blautia coccoides* (22), *Muribaculum intestinale* (20), *A. muciniphila* (14), and *Bifidobacterium animalis* (14) (**Fig. 2E**). However, *A. muciniphila*, not *B. caecimuris*, was by far the most abundant microbe in PedsCom mice after co-housing with OMM12 mice. Additionally, *B. coccoides* colonized poorly in PedsCom mice (~1%) despite ranking second in CAZymes missing in PedsCom. *Bifidobacterium intestinalis* and *M. intestinale* did not successfully colonize either OMM12 or PedsCom co-housed mice. *Enterocloster clostridioformis* ranked 7 out of 12 among OMM12 strains encoding CAZymes missing in PedsCom, yet it was the second most abundant strain in the cecum and third most abundant in the colon (**Fig. 1C and S1A**). Therefore, having quantitatively more CAZymes that are absent in PedsCom did not lead to higher OMM12 strain abundances in PedsCom mice.

The efficient colonization of PedsCom mice by the mucin-degrading specialist *A. muciniphila* suggested that foraging of host-derived glycans may contribute to microbial succession during weaning. In support of this hypothesis, the abundance of CAZymes that degrade host-derived glycans was strongly correlated with the relative abundance of the OMM12 strains in PedsCom mice co-housed with OMM12 mice (cecum, r=0.74, p=0.035; colon, r=0.75, p=0.031) (**Fig. 2F**). The total number of host-glycan-directed CAZymes present in a given strain also correlated with their relative abundances in the OMM12 control mice. Of note, we observed a non-significant trend between the abundance of CAZymes that metabolize fiber in each strain and their relative abundance in OMM12 control mice (**Fig S2B**). In contrast, the starch/oligosaccharide CAZymes are the only substrate group that is correlated with the relative abundances of the nine strains in PedsCom control mice (cecum, r=0.77, p=0.0015; colon, r=0.85, p=0.0034) (**Fig. S2C**). Ultimately, the total number of host-glycan degrading CAZymes best correlates with colonization success when introducing the adult-derived OMM12 strains into the PedsCom mice, indicating that host glycans provide an open niche in PedsCom mice that promotes microbial succession with microbes that can utilize host glycans such as mucus. Unexpectedly, host glycan utilization exceeds the advantage conferred by fiber-degrading CAZymes in other OMM12 taxa.

### Taxonomy predicts carbohydrate utilization profiles, but there are significant strain-specific differences in pre-weaning microbes

Though closely related lineages of microbial species often have similar preferences in carbohydrate utilization, any differences often indicate niche specialization^23–26^. At the lineage (taxonomical) level, we expected that the lower prevalence of fiber and host glycan CAZymes in PedsCom could be attributed to the higher proportion of *Bacilli* in this community, relative to adult-derived consortia that are composed primarily of *Clostridia* and *Bacteroidota*. Gut commensal *Bacteroidota* strains contain large numbers of CAZymes in their genomes^27^, and this difference is evident among the consortia strains (**Fig. S3A-B**) (median number of CAZyme per strain: *Bacteroidota* = 259, *Clostridia* = 102, *Bacilli* = 53, Other = 84). Non-metric multidimensional scaling (NMDS) clustering of CAZyme families from each strain by Bray-Curtis dissimilarity revealed clear separations between the *Bacteroidota, Clostridia* and *Bacillota* strains, emphasizing the impact of phylum level differences on metabolic capabilities (**Fig. 3A**), but strains from each community also occupied significantly different locations in NMDS space. Further, the PedsCom *Clostridia* were significantly more similar to *Bacilli* in CAZyme content than were the adult-derived *Clostridia* (**Fig. 3B**), with *Anaerostipes sp*. the most similar to *Bacilli*. These similarities suggested that the PedsCom Clostridia may possess metabolic capabilities more typical of *Bacilli* and less similar to canonical fiber-degrading *Clostridia* species.

**Figure 3.**
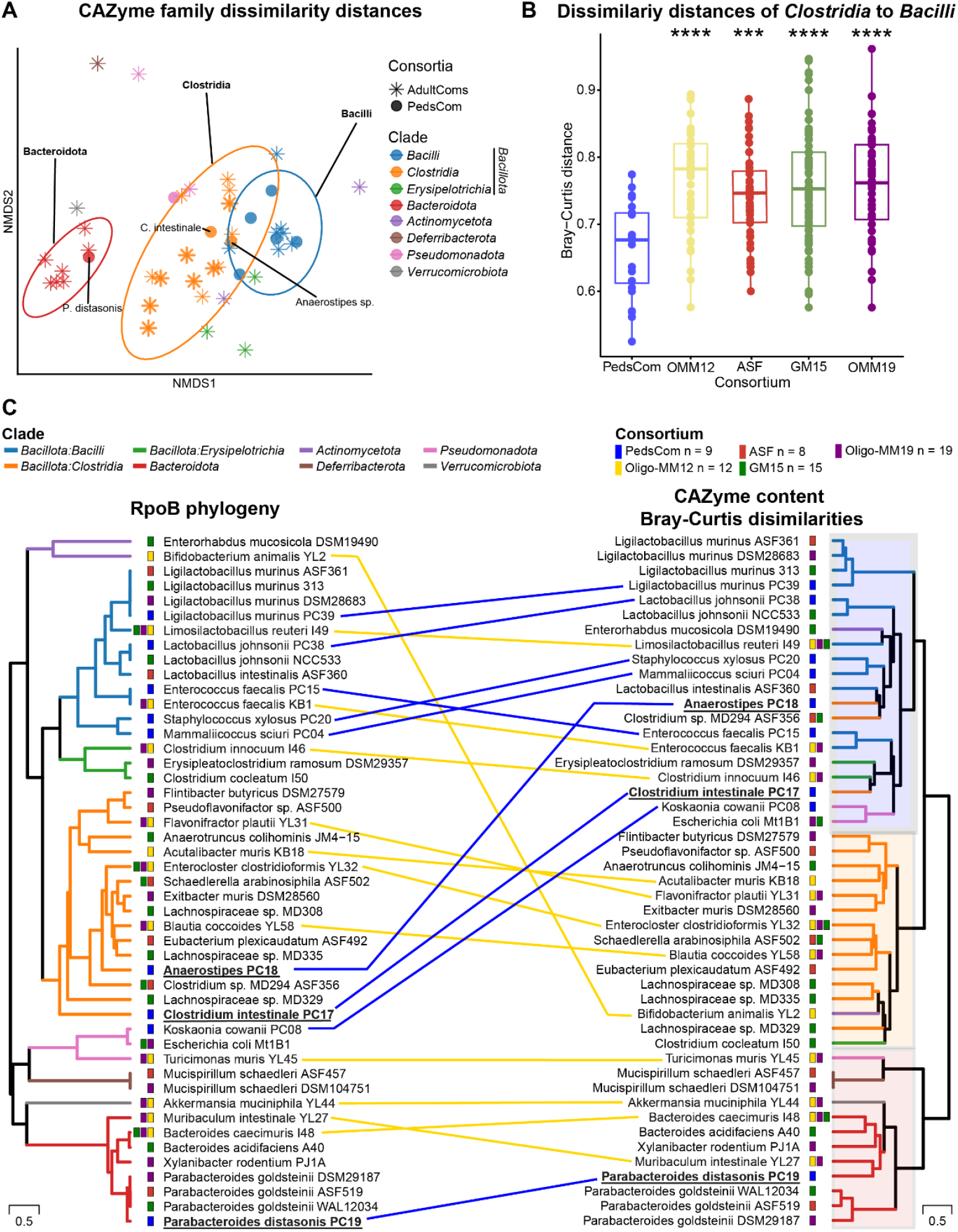
Taxonomy predicts carbohydrate utilization profiles, but there are significant strain-specific differences in pre-weaning microbes. **A**. Non-metric multidimensional scaling (NMDS) plot of Bray-Curtis dissimilarity distances of CAZyme families detected in consortia strains. Low variance (found in > 80% of strains) and rare (found in only one strain) CAZyme families were removed prior to analysis. 95% confidence intervals are shown for clades with 5+ strains (*Bacteroidota, Clostridia, Bacilli*). **B**. Box plots of the Bray-Curtis dissimilarity distances of consortia *Clostridia* (PC = 2, ASF = 5, GM15 = 6, OMM12 = 4, OMM19 = 5) to all *Bacilli* (12) strains. **C**. Tanglegram between the maximum-likelihood phylogeny of the RpoB amino acid sequence and hierarchical clustering based on Bray-Curtis dissimilarity of CAZyme families of consortia strains. The phylogenetic tree with the highest log likelihood is shown. Lines connect PedsCom (blue) and OMM12 (yellow) strain positions on each plot. Taxonomic clades are shown by branch color. Consortium membership denoted by colored bars near strain name. Shading on CAZyme dendrogram highlights distinct clusters of CAZyme family composition. Dunn’s test performed on pairwise consortia comparisons, False discovery rate (FDR) corrected, ***p<0.001, ****p<0.0001.

To determine how closely the phylogenetic relationships of the strains aligned with CAZyme content, we generated a phylogenetic tree of the five consortia using the highly conserved RNA polymerase subunit beta (RpoB) amino acid sequence. We used this tree to visualize the relationship between phylogeny and CAZyme content using a tanglegram with the hierarchical clustering of consortia strains by CAZyme family dissimilarities (**Fig. 3C**). As expected, the OMM12 strains of *Clostridia* clustered with the strains of *Clostridia* from the other adult-derived consortia (**Fig. 3C**). Unexpectedly, the PedsCom strains of *Clostridia, Anaerostipes sp*. and *Clostridium intestinale*, clustered with *Bacilli* in the CAZyme dendrogram. Additionally, *P. distasonis* possesses a distinct CAZyme profile compared to the adult-derived members of *Parabacteroides* genus, indicating a more similar CAZyme profile to phylogenetically more distant strains, e.g. the *Muribaculum* and *Xylanibacter* representatives (**Fig. 3C**). For most strains in these consortia the CAZyme content aligns well with phylogenetic grouping, but the *Clostridia* representatives from PedsCom share greater similarity to the carbohydrate metabolism of *Bacilli*, than the adult-derived *Clostridia*. Indeed, OMM12 *Clostridia*, on average, contain more fiber-degrading (18 vs. 7, PC vs OMM12) and host-glycan degrading (13 vs. 5, PC vs OMM12) CAZyme families than PedsCom *Clostridia*. This discrepancy between phylogeny and metabolic capabilities in the PedsCom *Clostridia* suggests that carbohydrate availability in the pre-weaning gut elicits selective pressure that influences which *Clostridia* strains colonize during the early-life period.

### Deletion of mucin transporter restricts *Akkermansia* colonization in PedsCom mice

To determine the extent to which glycan utilization is important for succession in PedsCom, we compared the efficiency of colonization of PedsCom mice with the type strain of *A. muciniphila* to an isogenic mutant that is unable to utilize mucus-derived host glycans due to a transposon insertion mutation of the mucin transport gene *mul1A* (amuc_0544)^28^ (**Fig. 4A**). The mean absolute abundance of wild-type *A. muciniphila* was ~100 times higher than the *mul1A* mutant (type strain = 1.1 ×10^11^ and mutant = 1.3 ×10^9^ *rpoB* copies/g feces) in PedsCom mice feces 1 day post gavage (**Fig. 4A**). Though the *mul1A* mutant was transiently able to reach wild-type levels 1 week post gavage, the *mul1A* mutant abundance was again significantly lower than the wild-type *A. muciniphila* at the 6-week time point. These data argue that mucus utilization supports efficient succession into the PedsCom community and is required to maintain a stable high-level colonization of *A. muciniphila* within this community.

**Figure 4.**
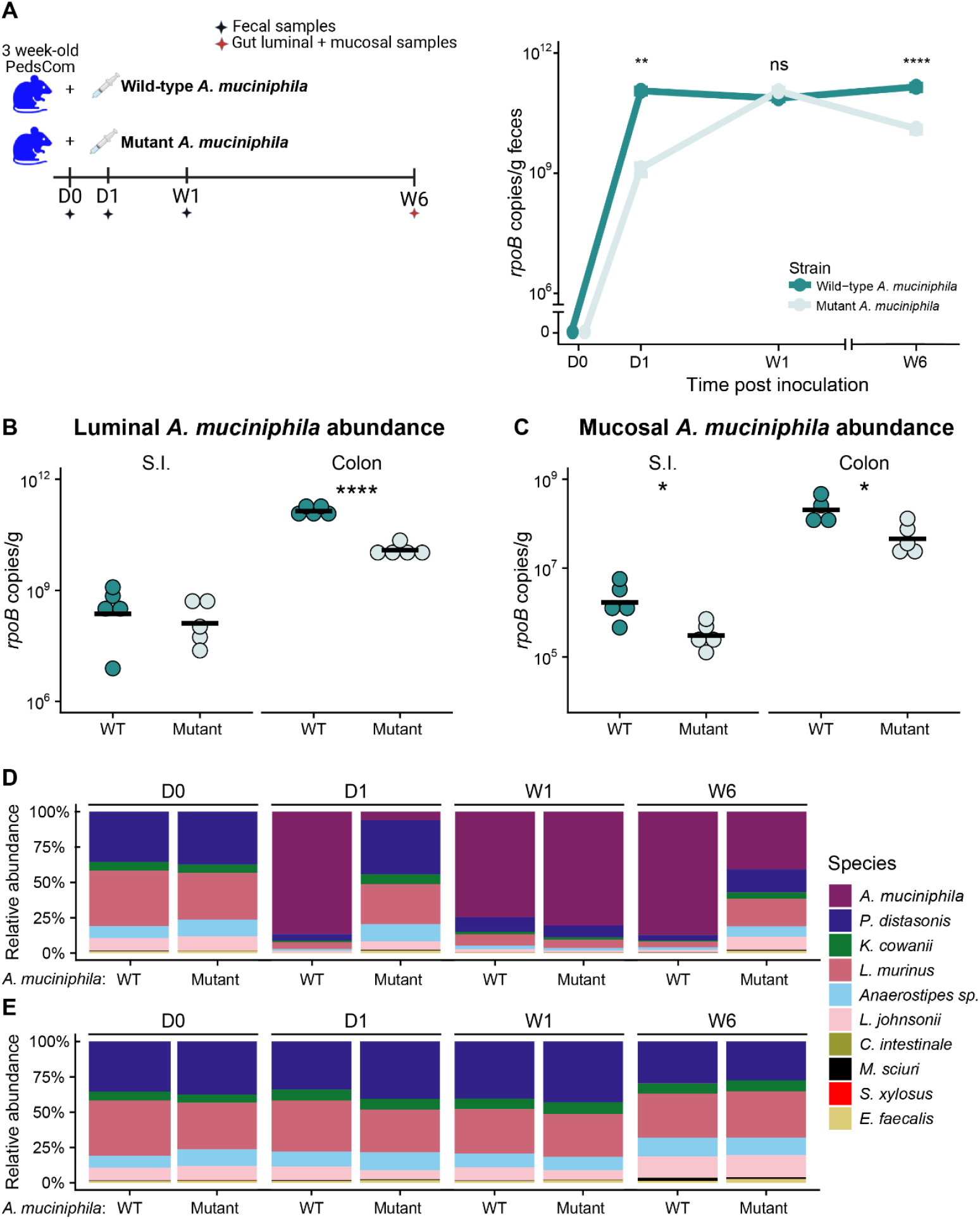
Deletion of mucin transporter restricts *Akkermansia* colonization in PedsCom mice. **A**. Schematic of *Akkermansia muciniphila* wild-type and *mul1A* mutant colonization experiment. Absolute abundance of PedsCom members after colonization with the wild-type (N = 5) or *mul1A* mutant (N = 5) strains of *A. muciniphila*. Abundance determined by *rpoB* copy number per gram/feces. **B**. Absolute abundances of wild-type (N = 5) and *mul1A* mutant (N = 5) colonization in PedsCom small intestine and colon luminal contents six weeks post gavage. **C**. Absolute abundances of wild-type [S.I. (N = 5), colon (N = 4)] and *mul1A* mutant (N = 5) colonization in PedsCom small intestine and colon mucosa six weeks post gavage. **D**. Mean relative abundances of *A. muciniphila* (wild-type or *mul1A* mutant) and PedsCom species in fecal samples. **E**. Mean relative abundances of PedsCom species in fecal samples, displayed without the relative abundance of *A. muciniphila*. Students t-test, *p<0.05, **p<0.01, ****p<0.0001.

To investigate the impact of mucus utilization on *A. muciniphila* colonization in different spatial compartments, we next assessed luminal and mucosal colonization of the wild-type and *mul1A* mutant *A. muciniphila* in PedsCom mice along the lower G.I. tract. The absolute abundance of the wild-type strain was significantly higher than the absolute abundance of *mul1A* mutant in the small intestine mucosa but similar in the small intestine luminal contents (**Fig. 4B-C**). In the colon, the *mul1A* mutant was less adept at colonizing the mucosa and lumen than wild-type *A. muciniphila*, with ~10 times lower abundances in both compartments. We observed patterns of colonization in the cecal mucosa and lumen similar to that of the colon (**Fig. S4A**). From these data, we conclude that the ability to utilize host-derived glycans from mucus provide a strong colonization advantage to *A. muciniphila* in the S.I., cecum and colon. The S.I. lumen is rich in dietary simple sugars, which likely accounts for the ability of the mutant *A. muciniphila* strain to colonize this nutrient-rich compartment at comparable levels to the wild-type strain. Overall, these *in vivo* data provide compelling evidence that the pre-weaning microbiome has an open niche for microbes that can utilize mucus glycans.

Since we proposed that *A. muciniphila* is filling an available host-glycan niche, we predicted that it would have minimal competition for resources with the other PedsCom microbes and have minimal impact on these communities. To address this hypothesis, we compared the relative abundance of PedsCom microbes after subtracting out the *A. muciniphila* reads. Strikingly, the introduction of wild-type or *mul1A* mutant *A. muciniphila* did not alter the relative or absolute abundances of the PedsCom consortia in the feces (**Fig. 4D-E, S4B)**, in the lumen or mucosa of the small intestines, cecum, and colon (**Fig. S4C-D**). However, there were small shifts in the abundance of the minor constituents of PedsCom community, based on the mucin utilization status of *A. muciniphila* (**Fig. S4E-F**). The overall lack of change in PedsCom abundance provides additional evidence that *A. muciniphila* occupies a distinct niche in the PedsCom mouse gastrointestinal tract and therefore does not compete with the most abundant PedsCom strains.

## Discussion

In this study, we developed a tractable *in vivo* approach combining gnotobiotic models of pre-weaning and adult microbial communities to investigate the microbial and host features that influence the weaning transition of the intestinal microbiome. Using this approach, we identified microbial degradation of host glycans as the microbial feature that best predicts which adult-associated microbes would effectively compete with pre-weaning microbes to colonize the gut. We validated the *in vivo* impact of host glycan metabolism by demonstrating that *A. muciniphila* requires mucus degradation capability to efficiently colonize PedsCom mice at weaning. Taken together, these gnotobiotic competition experiments between PedsCom and OMM12, *in silico* analysis of predicted metabolic functions of each strain in these communities, and *in vivo* studies of a strain of *A. muciniphila* harboring the mucus-degrading *mul1A* mutation, provide compelling evidence that the ability to degrade host glycans is a key driver of microbial succession around weaning.

The weaning transition to a solid-food diet is the largest nutritional change in the lifetime of mammals. This unique shift in nutrient availability leads to dramatic changes in the diversity, density, composition, and function of the intestinal microbiome over a relatively short time window (e.g., days for mice and a few months for humans). These remarkable changes in the microbiome drive critical features of immune system maturation and impact the risk of long-term immune dysregulation including colitis and autoimmunity^29,30^. Having a single dietary nutrient source during the pre-weaning period selects for microbes that are adept at utilizing these milk-derived nutrients. This intense nutritional selection is reflected in the highly distinct phylogenetic and metabolic profiles of the complex microbial communities of infants compared to adult communities^31^. Here we found that these distinctions are recapitulated in the gnotobiotic model community PedsCom compared to adult-derived microbial communities. PedsCom is dominated by *Bacilli*, such as *Lactobacillus*, and has few *Clostridia* species. The metabolic profiles of these *Clostridia, A. caccae* and *C. intestinale*, reveal them to be metabolically more similar to *Bacilli* than *Clostridia*, clearly illustrating the intense selection for microbes that are metabolically suited for the pre-weaning environment. As expected, the PedsCom consortium is metabolically adept at using the nutrients found in milk and is deficient in pathways involved in utilizing solid-food nutrients such as fiber. As the nutrient profile changes during weaning towards a fiber-rich, solid-food diet, the microbes that flourish under a milk-based diet face stiff competition from microbes that can utilize these new nutrients. Subsequently, the introduction of solid foods opens niches in the pre-weaning microbiome that can be exploited by adult-associated species to effectively compete with the existing pre-weaning communities to colonize, and in some cases dominate, the gut microbiome during the weaning transition.

While we expected PedsCom to be deficient in fiber degradation potential relative to adult-derived consortia, the deficiency in host glycan degradation was initially unexpected. Intestinal mucin production is the primary host-glycan that provides nutrients to commensal microbes in the gut of adult mammals^32^. Indeed, host glycan feeding of commensal microbes is a regulated process that helps maintain a healthy microbiome, particularly in times of stress such as starvation^33^. Intriguingly, mucin production, composition and degradation are developmentally regulated in mammals. For example, the expression of several mucin (*MUC)* genes and the degradation of intestinal mucus increase considerably during weaning^34–36^. The lower availability of mucins in neonatal mice likely leads to pre-weaning associated microbes being less reliant on mucin degradation as a nutrient source. *A. muciniphila* takes advantage of this available niche in a pre-weaning community to grow to high abundances in adult PedsCom mice. The *mul1A* mutation in *A. muciniphila* decreases colonization by ~100-fold, indicating the profound impact of mucus degradation on colonization efficiency for this important immunomodulatory microbe. It is noteworthy that *A. muciniphila* does not require this mucus degradation pathway to efficiently colonize germfree mice, indicating its ability to utilize other nutrient sources^28^, yet in the face of microbial competition, *A. muciniphila* relies on its ability to degrade and utilize the nutrients provided by the host mucins to compete in the mucosal and luminal niches of the gut. Similarly, the common gut commensal microbe *Bacteroides thetaiotaomicron* relies upon mucus utilization to successfully colonize the early-life microbiome before the introduction of plant-based fiber at weaning^45^. Taken together, the ability to utilize host-glycans is an important feature of commensal microbes that successfully colonize the early-life microbiome.

Mucins and HMOs are structurally similar, and this relationship potentially allows species that can use both nutrients to serve as transitional species during the weaning period^37^. One intriguing hypothesis is that the overlap of the metabolic genes that degrade and utilize the host-derived glycans mucin and milk oligosaccharides might help establish these early-life microbes in mucosal niches whose proximity to the host enhances the immunomodulatory potential of these microbes^38,39^. There are notable examples of gut commensals (*Bacteroides thetaiotaomicron, B. fragilis*, and *Bifidobacterium bifidum, B. longum*) that can utilize both mucins and human oligosaccharides (HMOs)^37,40,41^ and have well-described beneficial immunomodulatory effects^42,43^. *A. muciniphila* strains are also capable of metabolizing HMOs^44,45^, suggesting that *A. muciniphila* may also serve to bridge the gap between a pre-weaning and post-weaning microbiome. Furthermore, for bacteria that utilize mucin glycans but not HMOs, mucin degradation allows for persistence in the infant’s gut before the introduction of plant-derived fibers at weaning^46^. These findings implicate mucin-degradation as a significant mediator of microbial succession and immune system development during the weaning period.

Limitations: In this study, we developed a novel approach to model microbial succession of weaning using one defined pre-weaning community (PedsCom) and one adult-associated microbial community (OMM12). While these simplified communities recapitulate the features of pre-weaning and adult microbiomes^8,10^, it is possible that other defined communities, such as ASF or OMM19, may provide additional insights on metabolic features driving weaning succession. It is also likely that diets with different fiber content would yield additional insights into the interplay between fiber and host glycan degradation at weaning in promoting microbial and intestinal immune system maturation.

The weaning period is poised for probiotic interventions due to the low density and diversity of microbiota relative to adult communities, the emergence of new niches due to dietary changes, and several host-microbe interactions occurring during the weaning period that are crucial to long-term health^5^. Identifying the host and microbial features that drive succession at weaning is critical for developing effective therapies to shape healthy microbiome and immune system development. Our study suggests that the weaning period may be an ideal environment for colonization by microbes such as *A. muciniphila* that degrade host glycans. Understanding how carbohydrate sources affect microbiota succession will support the design of pre- and probiotic regimens to successfully colonize the infant gut with beneficial microbes, enhance immune development and improve long-term health.

## Methods

### Mice

Germfree and PedsCom colonized NOD mice were housed in flexible film isolators [Class Biologically Clean (CBClean), WI, USA] at the Hill Pavilion gnotobiotic mouse facility, University of Pennsylvania. Mice were fed LabDiet (IN, USA) 5021 (Cat# 0006540) and caged on autoclaved beta-chip hardwood bedding (Nepco, NY, USA). Animal studies were approved by the Institutional Animal Care and Use Committee (IACUC) of the University of Pennsylvania.

### PedsCom and Oligo-MM12 co-housing study

Oligo-MM12 strains were cultured and combined into an inoculum in sterile PBS+0.1% cysteine as previously described^8^. Oligo-MM12 inocula was orally gavaged into adult female germfree NOD mice twice, with three days between each dose, to generate the Oligo-MM12 gnotobiotic mouse line. Oligo-MM12 colonized mice were co-housed with adult female PedsCom mice for two days, then the Oligo-MM12 mice were removed and sacrificed. Co-housed PedsCom mice intestinal communities were allowed to stabilize for five weeks after which small intestine, cecum and colon samples were collected for metagenomic and gut lamina propria immune cell populations analyses.

### 16S rRNA gene metagenomic analysis

Metagenomic sequencing samples were stored at −80°C prior to extraction with the DNAeasy Powersoil kit, according to manufacturer instructions (Qiagen, Germany). The V4 variable region of the 16S rRNA gene was sequenced on the Illumina MiSeq platform by the PennCHOP microbiome program sequencing core as previously described^47^. Sequenced reads were processed into amplicon sequence variants using the QIIME2 pipeline (ver. 2023.7)^48^. Sequenced reads were denoised with Deblur (ver. 1.1.1)^49^. The SILVA 99% rRNA gene reference database (ver. 138) was used to assign taxonomic assignments to representative sequences. Reference sequences were further manually checked against the full length 16S rRNA gene sequence of each consortia strain using locally installed BLAST. Taxa bar plots were generated using the R package ggplot2 (ver. 3.5.1).

### Gut immune population analysis

Lamina propria immune cell populations were collected from mouse cecal and colonic tissues as previously described^50^. Splenocytes were collected and analyzed in parallel to serve as internal processing and staining controls. Cell surface markers were stained in RPMI, 4% FBS, in the dark, for 15 min on ice. Intracellular markers were stained by fixing and permeabilizing cell populations overnight at 4°C in permeabilization buffer (eBioscience, MA, USA) followed by staining in permeabilization buffer for 50 min, in the dark, at room temperature. The following fluorophore-conjugate antibodies and dilutions were used in this staining panel: BV510-CD45 1:200 (clone 30F11; Biolegend, CA, USA), PE-Cy7-TCRαβ 1:100 (clone H57-597; Biolegend), APC-Cy7-CD19 1:100 (clone 6D5; Biolegend), PerCP-Cy5.5-TCRγδ 1:100 (clone: UC7-13D5; Biolegend), AF700-CD8 1:100 (clone 53-6.7; Biolegend), FITC-CD4 1:100 (clone Rm4-5; Biolegend), APC-Foxp3 1:100 (FJK-16s; eBioscience), PE-RORγ 1:100 (clone B2D; eBioscience), Pacific Blue-Helios 1:33 (clone 22F6; Biolegend). Cell populations were analyzed on the LSRFortessa [Becton Dickinson (BD), MD, USA], and data analysis was performed on FlowJo v10.7 software (BD). Peripheral T regulatory cell populations were gated as: CD45^+^, TCRαβ^+^, CD4^+^, Foxp3^+^, RORγ^+^ and Helios^-^.

### Predicted carbohydrate metabolism analysis

Annotated proteomes of Oligo-MM12, ASF, GM15 and Oligo-MM19 species were downloaded from the Bacterial and Viral Bioinformatics Resource Center (BV-BRC) webserver. PedsCom species proteomes were annotated using the RAST tool kit (RASTtk) as previously described^8^. Predicted carbohydrate-active enzymes (CAZymes) in each species were identified using the dbCAN annotation tool ver.4.0.0^16^. CAZyme family designations were determined by successful annotation by at least two of the internal dbCAN predictive tools (HMMER, dbCAN_sub, Diamond). The metabolized substrates of each annotated CAZyme family were manually annotated using the CAZy database (https://www.cazy.org/) and dbCAN-sub database (https://bcb.unl.edu/dbCAN_sub/index.php). CAZyme family dissimilarity distances were computed using the Bray-Curtis method and visualized using non-metric multidimensional scaling with the R packages vegan (ver. 2.6-4) and ggplot2. CAZyme families found in only one (singletons) or over 80% of the consortium species were removed prior to analysis to limit uninformative functions. Hierarchical clustering (dendrogram) of consortium species based on CAZyme family dissimilarity was computed using square-root transformed Bray-Curtis dissimilarity matrix indices and the Ward D2 clustering method in base R.

### Phylogenetic analysis

The RNA polymerase subunit beta (RpoB) amino sequences of each consortium species were aligned with MAFFT (ver. 7.515) using the E-INS-I method^51^. The aligned sequences were used to generate a maximum likelihood tree using the L_Gasecuel_2008 amino acid substitution model with discrete gamma distribution (LG+G) with PhyML (ver. 3.3.20211231)^52^. Branch support was determined by Bayesian inference and the resulting tree was midpoint rooted. A tanglegram of the phylogenetic relationships of the consortium species and a dendrogram of CAZyme family dissimilarities was generated with the R packages ape (ver. 5.8) and dendextend (ver. 1.18.0) and untangled using the step2side method. Adobe Illustrator was used to label and highlight plots.

### Akkermansia muciniphila colonization

Type strain *A. muciniphila* BAA-835 was purchased through the American Type Culture Collection (VA, USA). A transposon insertion mutant in the mucin transport *mul1A* gene (amuc_0544) of *A. muciniphila* BAA-835 was previously generated^28^, and kindly provided to us by Raphael Valdivia.

*A. muciniphila* strains were grown under anerobic conditions (5% H_2_, 5% CO_2_, 90% N_2_) at 37°C in anaerobic *A. muciniphila* media supplemented with 0.5 g/L N-acetylglucosamine (Sigma, MO, USA)^53^. Three-week-old male PedsCom NOD mice were gavaged with 10^9^ CFUs of either wild-type or mutant *A. muciniphila*. Fecal samples were collected at day 0, 1 and 1 week-post gavage to assess PedsCom and *A. muciniphila* colonization dynamics. The mice were sacrificed at the six-week time point and small intestine, cecum and colon tissue and luminal contents were collected for analysis. PedsCom and *A. muciniphila* abundances were quantified by *rpoB* gene copies per gram of tissue (mucosal samples) or per gram of luminal contents (luminal samples) using quantitative PCR (qPCR). PedsCom species were quantified using previously described custom TaqMan primer and probe sets^8^. Primers were generated for this study targeting the *A. muciniphila rpoB* gene: forward – GTTCACGACCACATCGAAA, reverse – CGTCATCCAGTTCCGTAAAG, and analyzed using SYBR qPCR. Data acquisition was performed on the CFX Opus 96 PCR system (Bio-Rad, CA, USA) using TaqMan multiplex master mix (Thermofisher, MA, USA) and PowerUP SYBR Green PCR master mix (Thermofisher).

### Statistical analysis

Student T-test, Mann-Whitney, Kruskal-Wallis and Chi-squared analysis were performed using base R. Kruskal-Wallis post-hoc comparisons were performed by Dunn’s test with the Benjamini-Hochberg adjustment for multiple comparisons using the R package dunn.test (ver. 1.3.6). Chi-square post-hoc comparisons were performed with the Benjamini-Hochberg adjustment for multiple comparisons using the R package chisq.posthoc.test (ver. 0.1.2). Correlation plots were generated with Spearman’s rank correlation coefficients using the R package ggpubr (ver. 0.6.0.99).

## Acknowledgements

This study was supported by the National Institutes of Health T32 Postdoctoral Program in Clinical Pharmacology T32GM008562 (J.L.), National Institutes of Health grants R01DK133453-01A1 (M.A.S, P.J.P.), and JDRF grant 5-CDA-2020-946-S-B (M.A.S). We thank Dr. Adam Ratner for providing helpful discussion on this manuscript.

## Data availability statement

All data associated with this study is available upon request.

## Supplemental figure legends

**Figure S1. Relative abundances of PedsCom and Oligo-MM12 strains in co-housed mice**.

**A**. Individual bar plots of Oligo-MM12 strain relative abundance in Oligo-MM12 control, PedsCom control, Oligo-MM12 and PedsCom mice post co-housing. **B**. Individual bar plots of PedsCom strain relative abundance in Oligo-MM12 and PedsCom mice post co-housing. Bar plots represent median values.

**Figure S2. Host glycan CAZyme function predicts successful colonization of PedsCom mice**.

**A**. Bar plot of absolute count of annotated CAZymes in each consortium. **B**. Spearman correlations of CAZyme abundance, categorized by substrate, in Oligo-MM12 strains and relative abundance in Oligo-MM12 control (gold) and PedsCom co-housed mice (green) ceca and colons.

**C**. Spearman correlations of starch and oligosaccharide CAZyme abundance in PedsCom strains and relative abundance in PedsCom control mice ceca and colons. There were no significant correlations between CAZyme abundance and relative abundance of strains in the small intestine.

**Figure S3. *Bacteroidota* lineage is rich in CAZyme content**.

**A**. Bar plot of CAZyme count of each strain in this study ordered highest to lowest. **B**. Box plot of CAZyme count of PedsCom, Altered Schaedler Flora, GM15, Oligo-MM12, and Oligo-MM19 strains grouped by taxonomic clade.

**Figure S4. Defect in mucin utilization restricts *Akkermansia* colonization in PedsCom mice**.

**A**. Absolute abundances of wild-type (N = 5) and *mul1A* mutant (N = 5) colonization in PedsCom cecal luminal contents and mucosa six weeks post gavage. **B**. Absolute abundance of PedsCom species during following colonization by wild-type (N = 5) or *mul1A* mutant (N = 5) *A. muciniphila* fecal samples from PedsCom mice. Abundance represented by *rpoB* copy number per gram/feces. **C**. Mean relative abundances of *A. muciniphila* wild-type, *mul1A* mutant and PedsCom species in small intestine, cecum and colon six weeks post gavage. **D**. Mean fecal relative abundances of PedsCom species in small intestine, cecum and colon six weeks post gavage excluding *A. muciniphila*. **E**. Bar plots of the median relative abundance of *Clostridium intestinale* and *Enterococcus faecalis* in wild-type and *mul1A* mutant colonized PedsCom mice small intestine, cecum and colon luminal contents. **F**. Bar plots of the median relative abundance of *Clostridium intestinale* and *Enterococcus faecalis* in wild-type and *mul1A* mutant colonized PedsCom mice small intestine, cecum and colon mucosa. N = 5 for all conditions except for the wild-type mucosal samples (N = 4). Mann-Whitney test performed on median values, false discovery rate (FDR) corrected, *p<0.05.

