## Supplemental figures for "Microbial succession at weaning is guided by microbial metabolism of host glycans"

**Fig. S1 Relative of abundances of PedsCom and Oligo-MM12 isolates in co-housed mice**

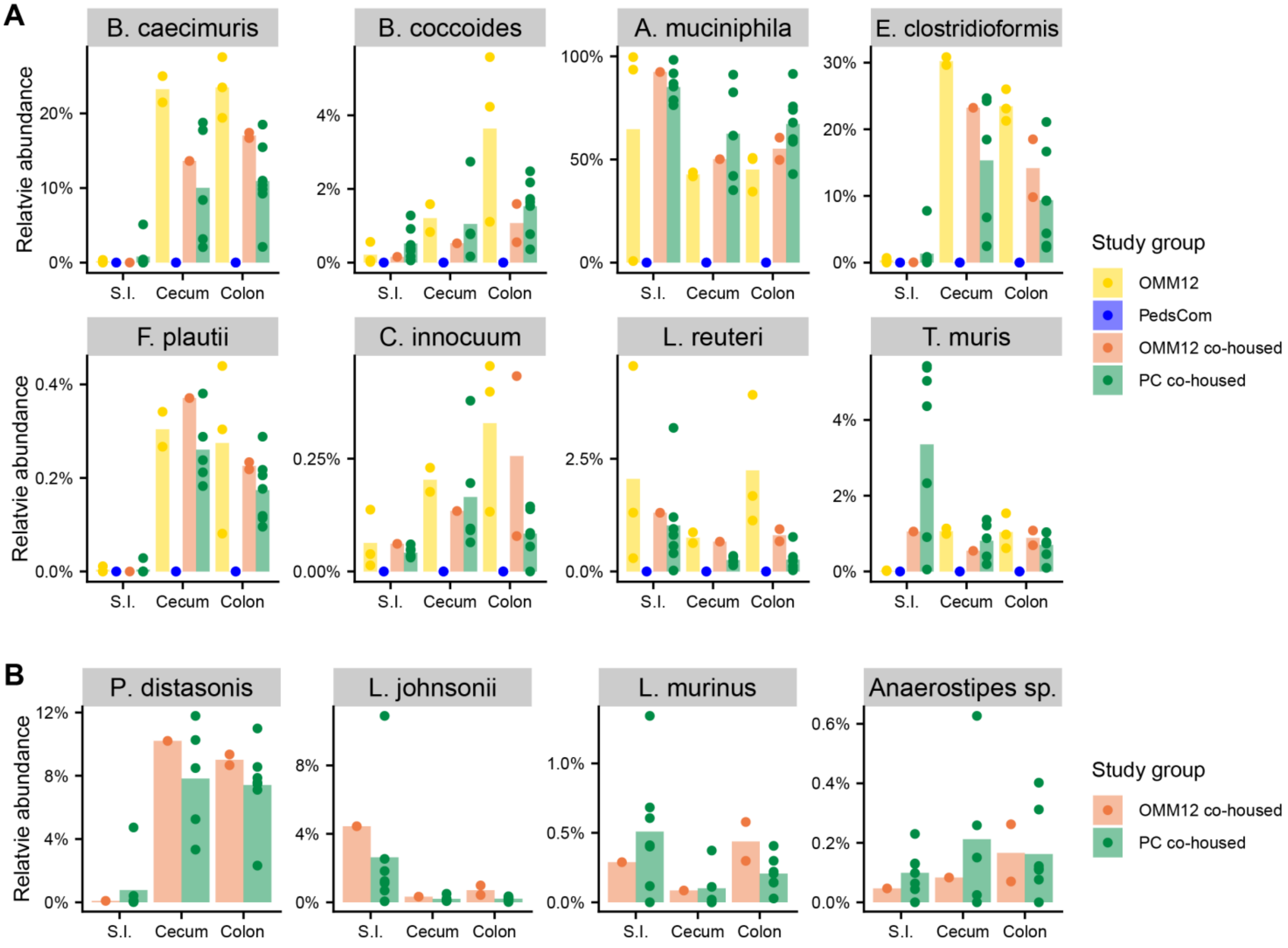

**Fig. S2 Host glycan CAZyme function predicts successful colonization of PedsCom mice**

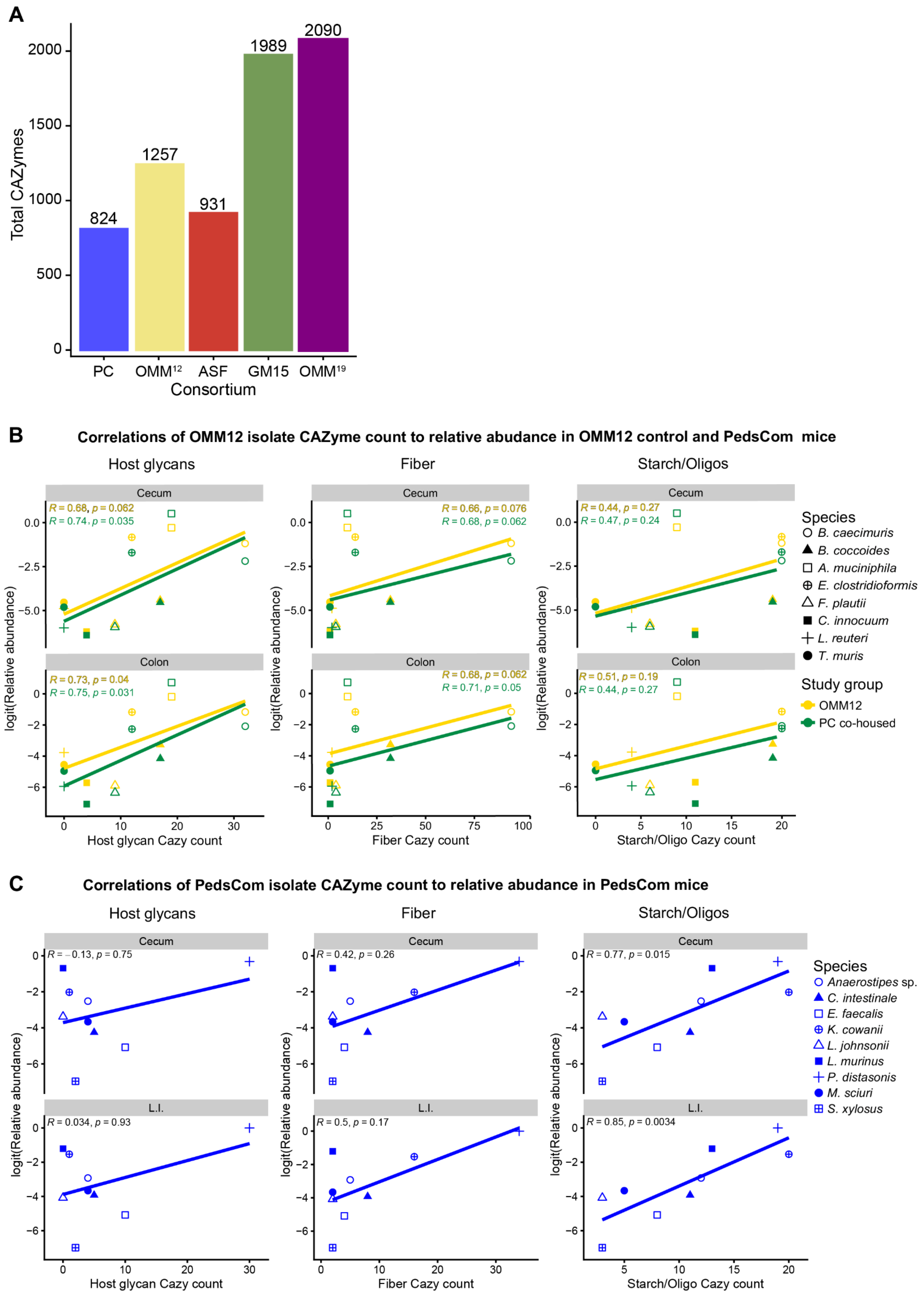

**Fig. S3 *Bacteroidota* lineage is rich in CAZyme content**

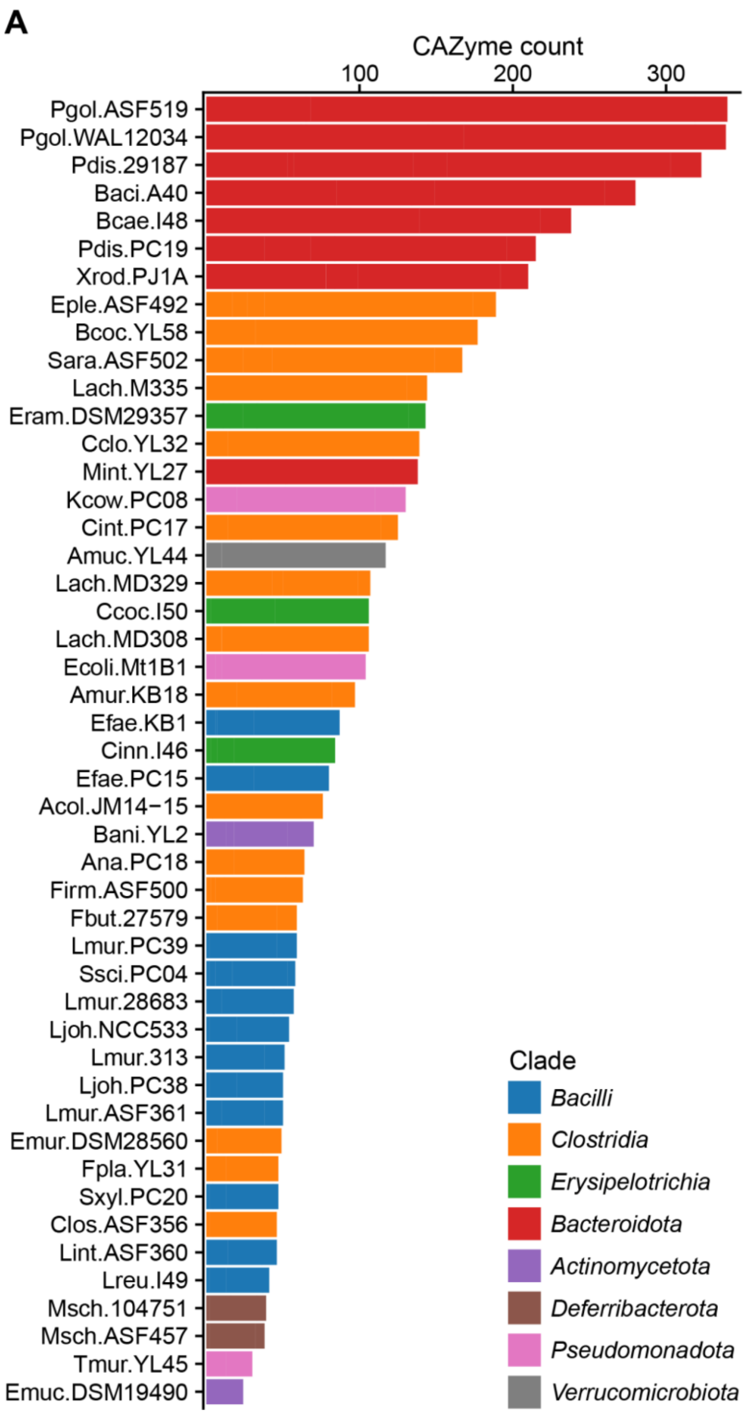

**Fig. S4 Defect in mucin utilization restricts *Akkermansia* colonization in PedsCom mice**

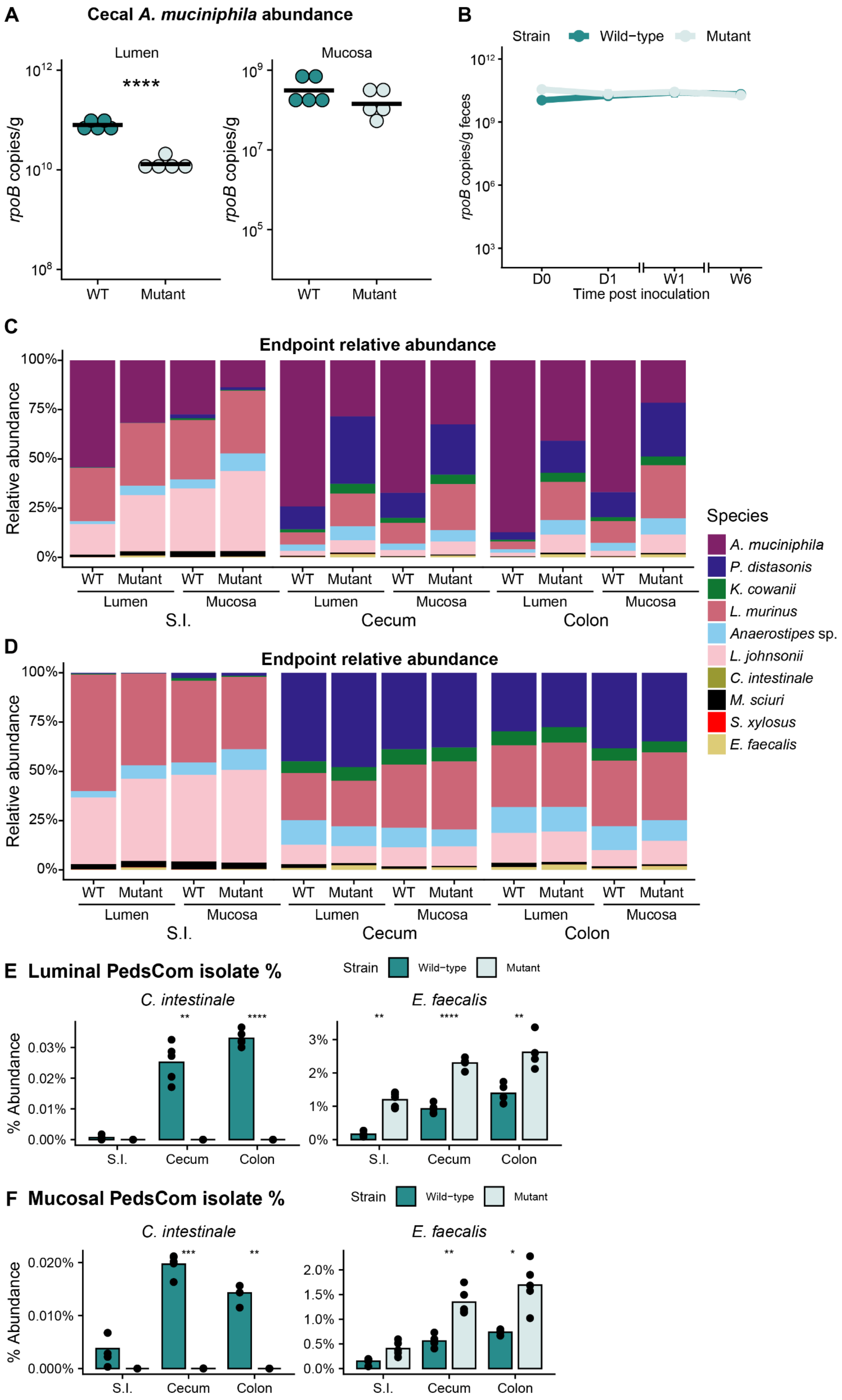
